# 3D reconstruction of the cerebellar germinal layer reveals intercytoplasmic connections between developing granule cells

**DOI:** 10.1101/2022.08.21.504684

**Authors:** Diégo Cordero Cervantes, Harshavardhan Khare, Alyssa Michelle Wilson, Nathaly Dongo Mendoza, Orfane Coulon--Mahdi, Jeff William Lichtman, Chiara Zurzolo

## Abstract

**Summary:** The difficulty of retrieving high-resolution, *in vivo* evidence of the proliferative- and migratory processes occurring in neural germinal zones has limited our understanding of neurodevelopmental mechanisms. Here, we employed a connectomic approach using a high-resolution, serial-sectioning scanning electron microscopy volume to investigate the laminar cytoarchitecture of the transient external granular layer (EGL) of the developing cerebellum, where granule cells coordinate a series of mitotic and migratory events. By integrating image segmentation, 3D reconstruction, and deep learning approaches, we discovered and characterized anatomically complex intercellular connections bridging pairs of cerebellar granule cells throughout the EGL. Connected cells were either mitotic, migratory, or transitioning between these two cell stages, displaying a chronological continuum of proliferative and migratory events never previously observed *in vivo* at this resolution. This unprecedented ultra-structural characterization poses intriguing hypotheses about intercellular connectivity between developing progenitors, and its possible role in the development of the central nervous system (CNS).

## Introduction

The principles that regulate the development of brain connectivity and modulation of synaptic transmission are just beginning to be understood (Witvliet et al., 2021; Yuste et al., 1995). From the historical controversy between the reticulon and neuron theories (https://www.nobelprize.org/prizes/medicine/1906/golgi/lecture/, https://www.nobelprize.org/prizes/medicine/1906/cajal/lecture/) it is plausible that synaptic transmission does not solely explain how brain cells communicate.

During development, brain cells communicate in order to orchestrate cell divisions and migrations in the absence of synapses;, this intercellular communication relies significantly on gap junctions and mitogens (Wang et al., 2022) which contribute to the establishment of neuronal assemblies.

Another mechanism of intercellular communication allowing the exchange of molecules as well as entire organelles between connected cells are stable intercellular bridges (IBs), which arise from incomplete cytokinesis – part of the cell division process during which the cytoplasm of a single cell is stalled as it divides into two daughter cells. Unlike cytokinetic bridges (CBs) that form and are normally cleaved in a series of well-known events during cytokinesis (Green et al., 2012), the mechanisms regulating the development of IBs and their function are not yet understood. Today, IBs are considered an evolutionarily conserved process across species (from insects to humans) that occur in female and male germ lines; they have rarely been observed in somatic tissues of invertebrates and never previously in the brain (Haglund et al., 2011).

The cerebellum – a brain region vital for balance control and motor movements – arises from a series of finely-orchestrated cellular division and migratory events. These processes are paramount for proper development of the cerebellar cortex, and in particular, the postnatal development of granule cell (GC) progenitors within the transient, germinal External Granular Layer (EGL) (Cajal, S.R., 1888).

GCs are thought to transition through symmetric and asymmetric divisions exclusively in the outer-EGL (oEGL) sublayer before gradually exiting the cell cycle and differentiating into migratory, post-mitotic GC neurons in the inner-EGL (iEGL) (Miyashita et al., 2017). Although previous studies have focused on the molecular pathways regulating the exit of GCs from the EGL (Famulski et al., 2010), the lack of a morphological survey of GCs has hindered a detailed description of the postnatally developing cerebellar cortex.

To shed light on this gap, we examined the EGL of a developing mouse cerebellum (lobule VIII) using a serial-sectioning scanning electron microscopy- (ssSEM) based connectomic approach, which represents a state-of-the-art approach to study brain connectivity at nanometer resolution (Kasthuri et al., 2015; Wilson et al., 2019). Obtained from a 7-day-old mouse (postnatal day 7 or P7), this dataset exhibited an expanded EGL resulting from a peak in GC proliferation at P6 (Espinosa and Luo, 2008) that enabled us to study a large number of GCs in the EGL. These data led to the discovery of GCs connected by membrane-bound cytoplasmic, intercellular connections (ICs) throughout the depth of the EGL. We analyzed the localization, morphology, and subcellular architecture of connected GCs to investigate their origin. Whereas some of these connections exhibited similarities with (CBs), most displayed features that were never previously described, and could be unique to the brain.

In summary, our systematic examination of the neural germinal layer of the cerebellar cortex by ssSEM revealed frequent cytoplasmic interconnections between developing neural cells. Given persistent CBs have never been observed in the brain, our work highlights unknown morphological and cytological features of developing GCs in the mouse cerebellum and provides a new perspective on the mechanisms regulating brain development as a whole.

## Results

### Identification of cerebellar granule cells linked by intercellular connections using ssSEM

We obtained a ssSEM volume spanning 1.7×10^6^ μm^3^ in volume at a voxel size of 4×4×30 nm^3^ (Wilson et al., 2019) from the developing cerebellum of a P7 mouse (lobule VIII) (**Fig. 1A-B, Movie S1)**. To analyze the morphological features of GCs in the EGL, we manually segmented the soma and protrusions of cells in the volume using *VAST* (Berger et al., 2018). This led to the striking observation of a pair of GCs linked by a membranous, IC resembling a CB that forms towards the end of mitosis (**Fig. 1C-E’; Movie S2**). CBs are microtubule-rich connections that play a vital role in the abscission of dividing cells; however, observations of CBs have been limited to *in vitro* models (Andrade et al., 2022; Guizetti et al., 2011). Full segmentation of an IC revealed two features previously ascribed to CBs: (1) an electron-dense structure within the tube, which could indicate the presence of a midbody (MB); and (2) microtubules emanating from the dense region into both cells (**Fig. 1D-E’**) (Sherman et al., 2016).

**Fig. 1.**
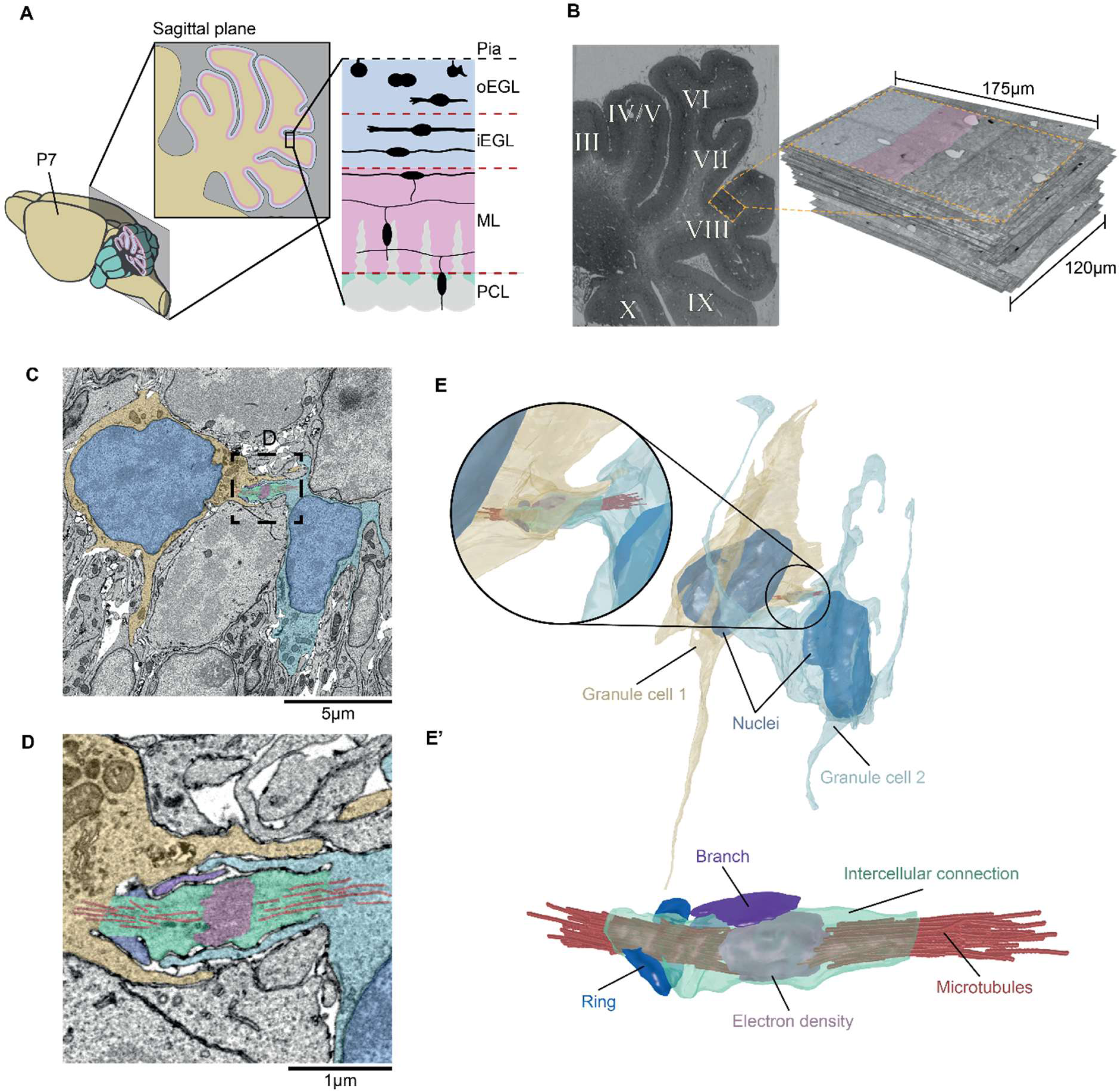
Identification of GC intercellular connectivity. **(A)** Schematic of P7 mouse cerebellar cortex sagittal cross-section. Sagittal view of cerebellar lobule depicts external granular layer (EGL) divided into outer & inner EGL (oEGL; iEGL). During early stages of post-natal development, GC progenitors populate only the oEGL where they proliferate through mitosis. The postmitotic GCs arising from these divisions then migrate in the iEGL, performing tangential migration. In order to become mature neurons, the postmitotic GCs then undergo radial migration through the Molecular Layer (ML) and the Purkinje Cell Layer (PCL). The ML is located underneath the EGL, and is mostly made of the parallel fibers extended by the GCs doing radial migration. GCs’ journey ends when they settle in the Internal Granular Layer (IGL), where they establish synapses with Golgi cells and Mossy fibers (not shown). (**B**) Electron micrograph (left) and 3D volume of P7 mouse, lobule VIII (right) prepared using serial sectioning scanning electron microscopy (ssSEM) (blue, oEGL; pink, iEGL. (**C-D**) 2D electron micrographs showing GCs in the EGL bridged by an intercellular connection (IC). (**E**-**E’**) 3D reconstructions of (**C)** and **(D**), respectively, showing microtubules (red) that emanate into both cells, an electron-dense region at the center of the IC (pink), and a branch (magenta) and an incomplete ring (blue) extruding from the IC (green).

Connections between GCs in the EGL were not previously reported; thus, to quantify and characterize this phenomenon, we executed a systematic screen of all ICs in the EGL of our volume. To focus on GCs of the EGL, and not on cells migrating radially through the molecular layer (ML) (**Fig. 1A**), the neighboring region which also contains other interneurons, we first determined the EGL|ML boundary through an automatic and reproducible approach. We accomplished this by designing our own 2D Convolutional Neural Network (CNN) architecture that was iteratively applied to the slices of our volume, dramatically reducing the manual effort and time required to distinguish the two layers (**See supplementary methods, Fig. 2A and S1**).

**Fig. 2.**
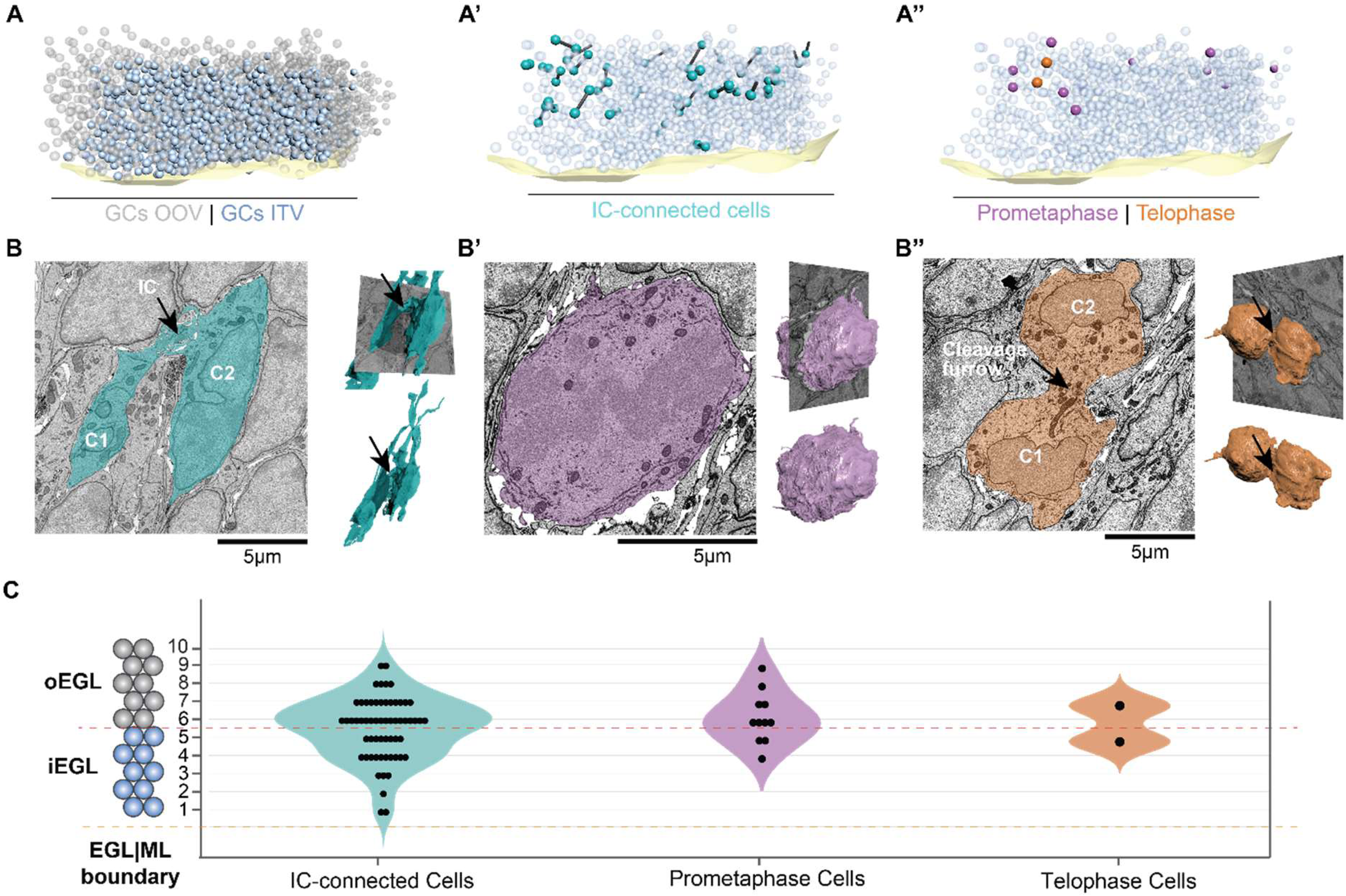
Characterization of GCs in the ssSEM volume. **(A)** 3D map of EGL|ML boundary (light yellow) and 2042 GCs in the EGL of the volume: 1110 partially outside of the volume (OOV-gray); 932 inside the volume (ITV-blue). **(A’)** 3D map of IC-connected GCs (cells-cyan, ICs-black cylinders). **(A’’)** 11 GCs in prometaphase (purple) and 2 in telophase (orange) distributed ITV. **(B, B’ and B”)** 2D cross-sections along the sagittal plane of the EGL (left) and 3D reconstructions (right) of GCs in different cell cycle stages. **(B)** IC-connected GCs (black arrow) with well-formed nuclei and long protrusions; **(B’)** GC in prometaphase exhibiting a broken nuclear envelope and a round shape without protrusions; **(B’’)** GCs in telophase showing fully-formed nuclei, without protrusions, and connected by a cleavage furrow (black arrow). **(C)** Distribution of GCs interconnected by ICs (n=60), prometaphase (n=11), and telophase (n=2) throughout the EGL shows that all events take place in the inner (layers 1-5) and outer (layers 6-10) EGL.

### GCs carry out mitotic divisions across the EGL

We manually identified and counted all GCs in the volume **(See supplementary methods)** and identified a total of 2042 GCs. Of these, 1110 cell somas were partially outside the volume (OTV) as they were located at the edges of the volume. We focused our study on the 932 cells with nuclei inside the volume (ITV) (**Fig. 2A**).

A subcellular diagnosis of every ssSEM slice occupied by each GC revealed 60/932 GCs forming ICs (6.4% of cells) (**Fig. 2A’ and 2B**). In addition, we also identified 11/932 GCs in prometaphase (1.2% of cells), evidenced by a disassembled nuclear envelope (Beaudouin et al., 2002) (**Fig. 2A’’ and 2B’**); and 2/932 GCs in telophase (0.2% of cells), identified based on their spherical-shaped morphology, assembled nuclei; and wide, cleavage furrow (**Fig. 2A’’ and 2B’’**).

This observation prompted us to investigate if the laminar location of these three distinct cellular events was consistent with previous reports of GCs exclusively dividing in the EGL, as well as to test the proximity of ICs in relation to dividing GCs. To this end, we split the EGL into two constituent, 5-cell-deep sub-layers representing the oEGL and iEGL, as previously reported in the literature (Cadilhac et al., 2021) (**Fig. 2C**). Layer numbers were manually assigned to every GC by counting the number of cells between the ML and the GC of interest. Using this definition, 25 (42 %) of IC-connected GCs belonged to the oEGL and 35 (58%) to the iEGL. On the other hand, 8 (73%) of the prometaphase cells belonged to the oEGL and 3 (37%) to the iEGL (**Fig. 2C**). Telophase cells were located at the boundary between the o- and iEGL **(Fig. 2C)**. These latter observations indicate that GCs don’t only divide in the oEGL as previously thought (Gao and Hatten, 1993), but also in the iEGL (Hanzel et al., 2019). Furthermore IC-connected cells significantly outnumbered GCs in prometaphase and telophase, challenging the possibility that all ICs are mitosis-derived CBs. We therefore decided to study the morphological features of IC-connected cells and explored whether these characteristics could shed light on their origin and function.

### Morphological features of connected GCs demonstrate structural differences between ICs and CBs

By using 3D reconstructions of fully segmented IC-connected cells, we revealed three intriguing morphological features that may differentiate ICs from classical CBs.

First, in contrast to the stereotypical CB that connects the soma of two cells and aligns predominantly with the nuclei of daughter cells (Andrade et al., 2022), some of our ICs connected to other parts of the cell, at times even connecting a process of one cell to the soma of the other, rather than simply connecting soma to soma in an aligned fashion (**Fig. 3A**).

**Fig. 3.**
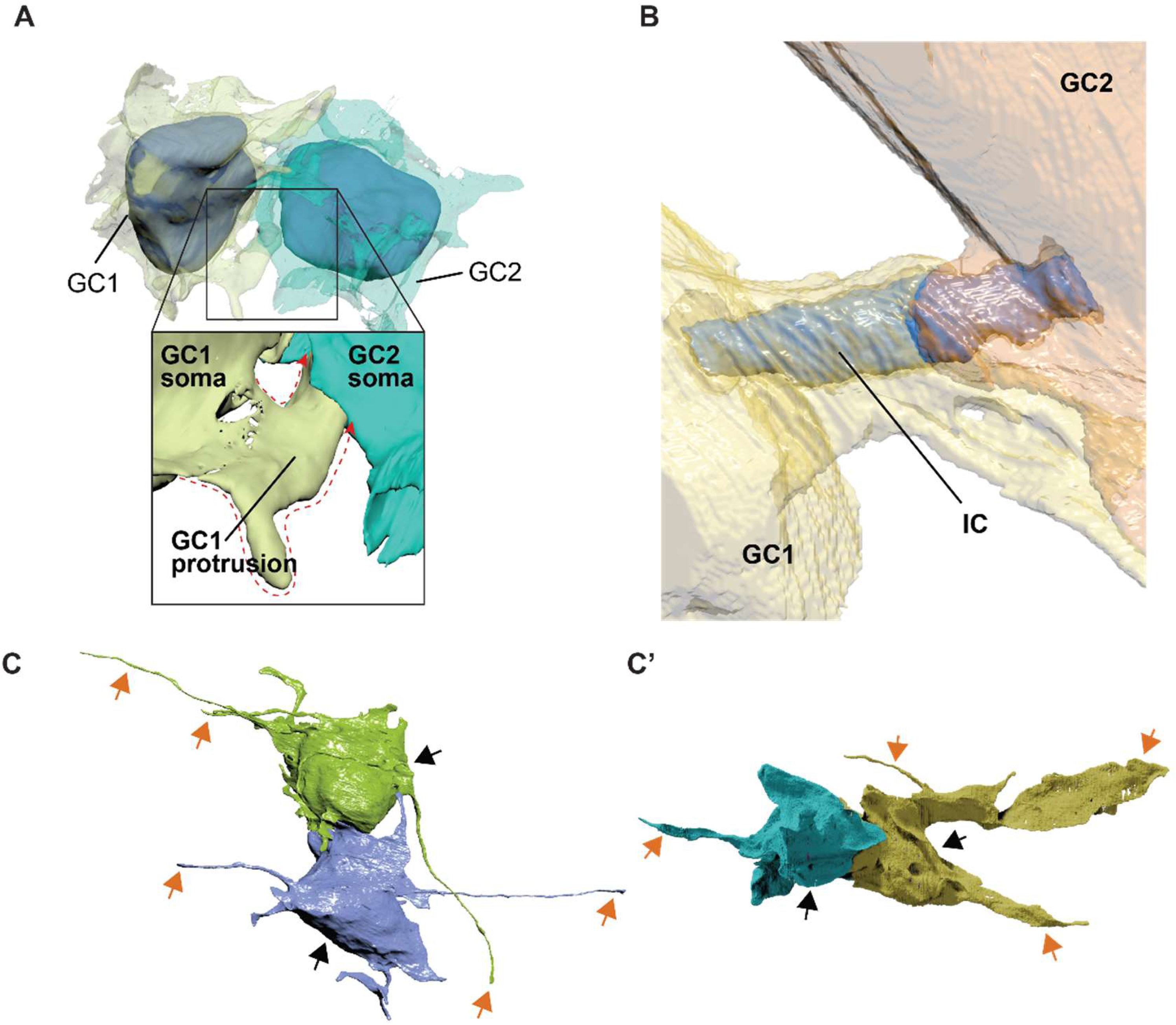
Morphological assessment of IC-connected GCs reveals distinct cellular features. (**A**) 3D reconstruction showing an example of IC-connected GCs via a cell protrusion (protrusion contour depicted by red dotted line): GC1 and GC2 connected via IC originating from protrusion of GC1. (**B**) 3D reconstruction showing sheets wrapped around IC (blue) extending from both connected cells (GC1, yellow; GC2, orange). (**C, C’**) 3D reconstruction of IC-connected GCs (cell somas-black arrows) extending long protrusions (orange arrows) as filopodia (left) and lamellipodia (right).

Second, ICs were not always directly exposed to the extracellular matrix, as is the case for CBs *in vitro* (Sherman et al., 2016) and *in vivo* (Rathbun et al., 2020). Instead, they were wrapped by membranous lamellipodia-like sheets that stemmed from both connected cells (**Fig. 3B**).

Third, in several cases, IC-connected GCs extended protrusions of up to 31.15 μm in length **(Fig. 3C and 3C”**), which is an unexpected behavior of dividing GCs as membranous processes are thought to develop after cells disconnect from each other post cytokinesis (Hanzel et al., 2019). Although GCs are expected to have extended parallel fibers by the end of their tangential migration, GCs with protrusions being connected makes this observation unique.

### IC-connected cells are mitotic, migrating, or transitioning between the two states

To determine whether ICs could have resulted from cell division, we investigated the cell-cycle stages of IC-connected GCs. This was inferred morphologically, both by 3D-reconstruction of cell shape and by characterization of subcellular features that change significantly during different stages of the cell cycle. These included the number and distribution of Golgi complexes; the position of centrosomes; and the presence of a primary cilium (**Fig. 4A-C, S2)**. We selected 21 IC-connected GC pairs for this analysis (i.e. 42 GCs), specifically because all morphological features in consideration were visible in the volume.

**Fig. 4.**
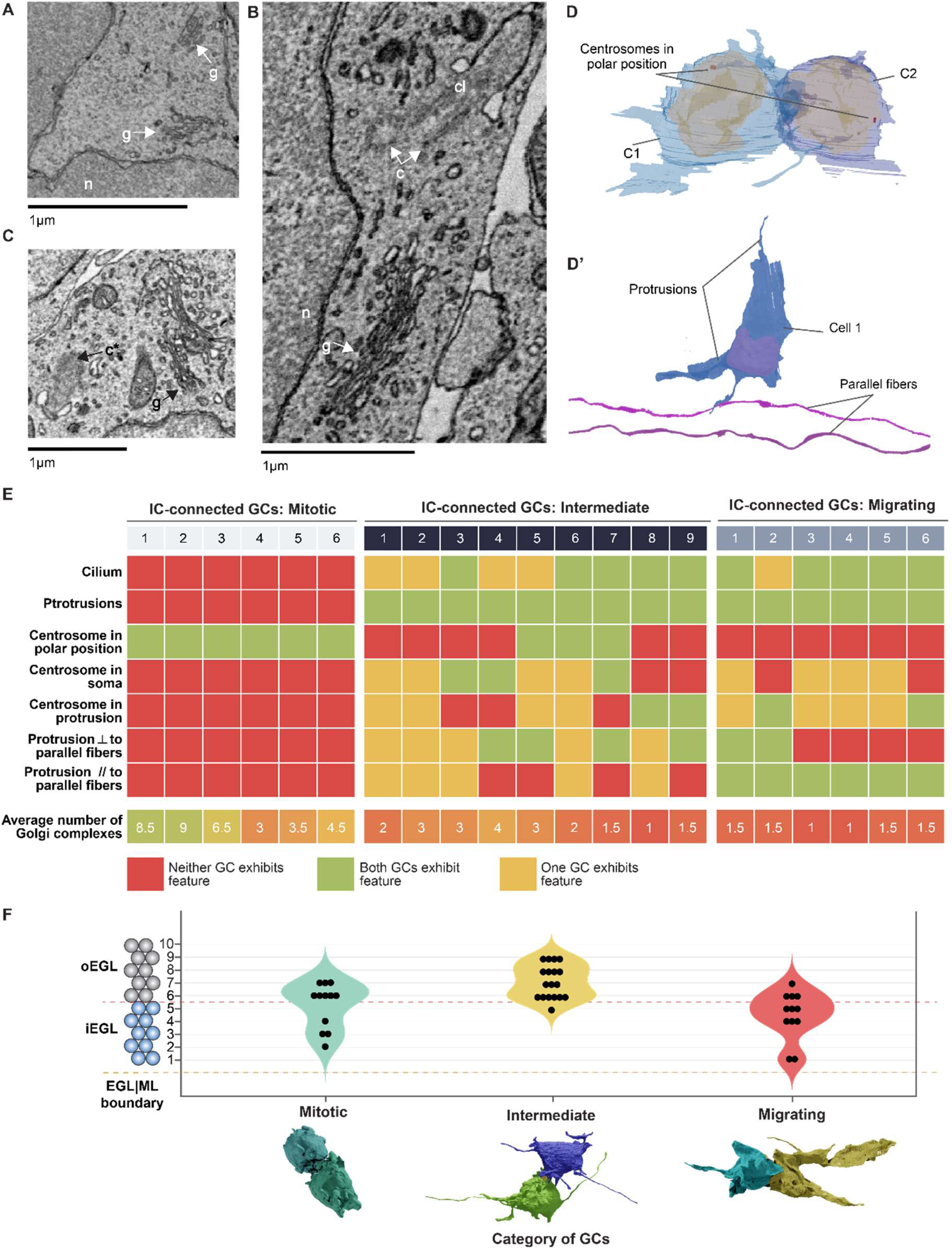
IC-connected cells are mitotic, migrating, or transitioning between the two states (intermediate) **(A, B, C)** Identification of cellular organelles within GCs in electron micrographs: n-nucleus, c-centrosome, g-Golgi complex, c*-centriole forming a cilium, cl-cilium. **(D)** IC-connected GCs showing centrosomes at polar positions indicating a mitotic stage of GCs. **(D’)** GC showing protrusions and two parallel fibers (purple) as a reference for orientation; the protrusion orientation was described with respect to the parallel fibers. **(E)** Summary of features based on cellular organelles and protrusions for three categories of IC-connected GCs: mitotic, intermediate and migrating. Total 42 GCs (21 ICs) considered for this analysis. * ┴ stands for perpendicular, **//** stands for parallel **(F)** The distribution of IC-connected GCs (n=42) in three categories: Mitotic (n=12), intermediate (n=18) and migrating (n=12).

Our analysis revealed three major categories of IC-connected GC-pairs: mitotic, migrating, and intermediate **(Fig. S2)**. Of these, 6 GC pairs (12 GCs) were mitotic. These cells had more than two Golgi complexes dispersed in the cell (**Fig. 4E, and S3A, S3B)** and were characterized by the absence of a cilium, both features of dividing cells (Colanzi and Corda, 2007; Frye et al., 2020; Plotnikova et al., 2009). Accordingly, these cells were spheroid in shape (**Fig. 4D, S2)**, and had centrosomes positioned at opposite cell poles, probably corresponding to spindle poles (Tang and Marshall, 2012). Altogether, these observations suggest that these cells may be at the end of mitosis, presumably in late telophase/early cytokinesis (Schweitzer et al., 2005).

Another 6 pairs (12 GCs) were migrating. These cells harbored fewer than 2 Golgi complexes and bore a cilium, indicating a G0/G1 stage (**Fig. 4E, S3A, S3B**). They also showed well-developed lamellipodia oriented parallel to the EGL|ML boundary and parallel fibers, a characteristic of tangential migration (Komuro and Yacubova, 2003) (**Fig. 1A, S2**). Most of these GC pairs were found in the iEGL, consistent with their tangentially migrating stage.

Based on the morphological criteria analyzed above, we could not categorize the remaining 9 pairs as undergoing migration or cell division, and were therefore defined as ‘intermediate’ (**Fig. 4E**). These cells were not “truly” migrating tangentially as their protrusions did not show any specific orientation (**Fig. 4D’, S2**) and they were mostly located in the oEGL (**Fig. 4F**). These GCs exhibited diverse shapes-for instance, one pair showed cells with a round soma and long filopodia, while two other pairs had elongated somas and small lamellipodia. Intermediate cells also shared morphological features with mitotic and migrating cells. For example, some had more than two Golgi complexes and a cilium. In these cells, the centrosome was mainly located in the soma (not close to a protrusion), supporting the fact that they were not undergoing migration. Thus, this analysis suggests an overlap of events at the end of mitosis and extension of protrusions (**Fig. 4E**).

Interestingly, 2 mitotic GC pairs were located deep in the iEGL (**Fig. 4F)**, reinforcing the observation **(Fig. 2C)** that cell division doesn’t only occur in the oEGL as previously proposed (Gao and Hatten, 1993), but also in the iEGL. Similarly, 4 of 12 migrating GCs were observed in the oEGL, implying that migration is not restricted to the iEGL (**Fig. 4F)**.

In 8 of 12 migrating GCs, the centrosome was located close to one of the protrusions, as expected for migrating neurons, where centrosomes have been shown to be in the vicinity of the leading process or a lamellipodium (Bellion et al., 2005; Tsai and Gleeson, 2005). For the remaining 4 migrating GCs, the centrosome was in the soma, and for 2 of these GCs it was also close to the IC. These observations could imply that the centrosome was no longer in charge of spindle poles and GCs in the migrating category have exited mitosis.

### ICs are anatomically diverse and complex

In order to characterize the diversity of ICs, we developed a program for morphometric analysis of segmented cells called *CellWalker* that allows for calculation of morphological features of segmented cellular structures, including length, diameter, volume, surface area and curvature **(see supplementary methods)**. Using CellWalker, we saw that IC lengths ranged from 1.25-to 3.3 μm with an average of 2.25 μm (**Fig. 5A**), comparable to *in vitro* and *in vivo* CBs, which exhibit lengths between 3 and 5 μm (Mullins and McIntosh, 1982).

**Fig. 5.**
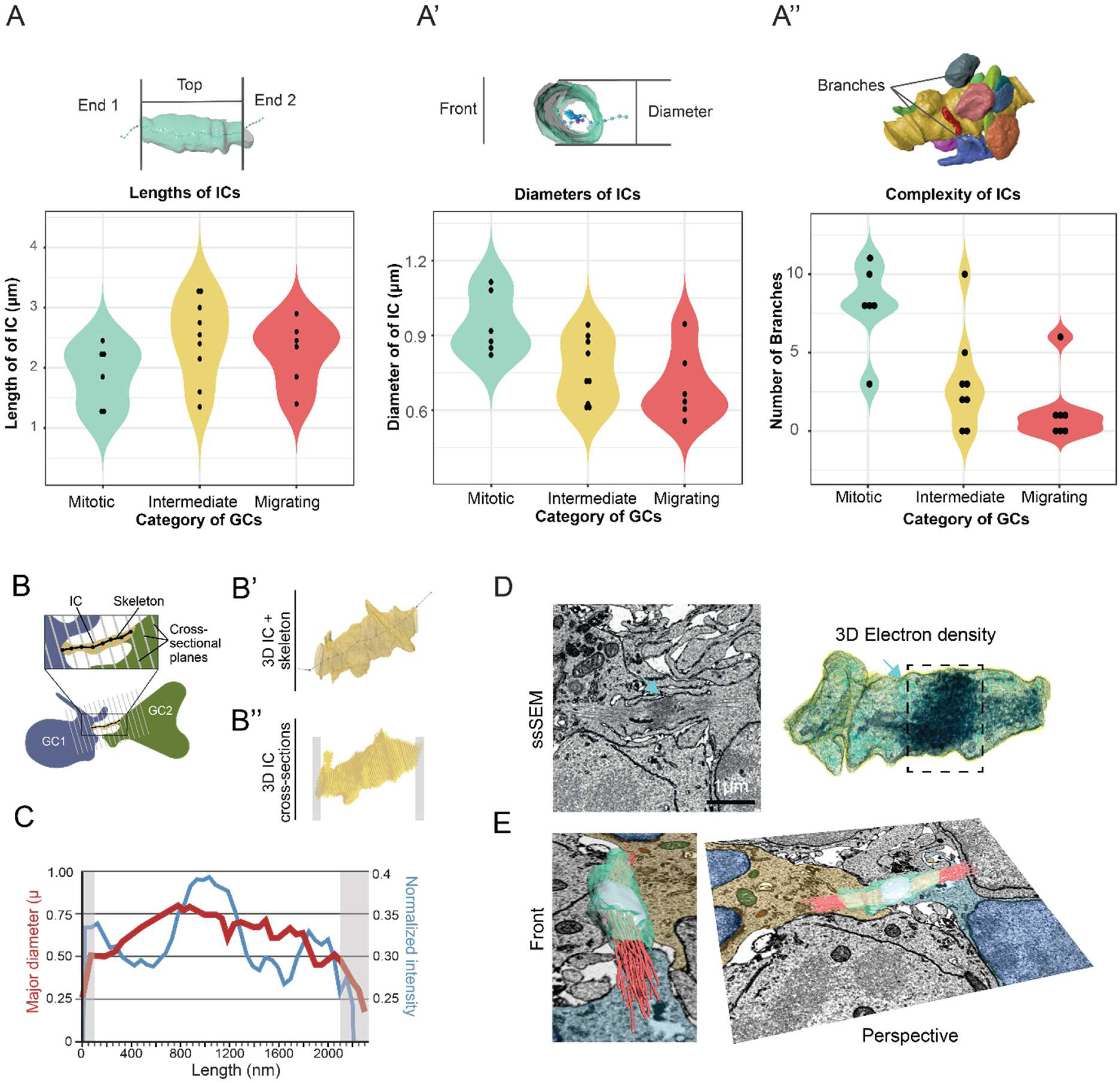
Morphologic characterization of the connection in interconnected GCs. (**A**) 3D reconstruction of an IC with a skeleton. End 1 and End 2 mark sites of contact with GCs, depicting the length calculation of ICs. **(A’)** IC with a skeleton from front view showing diameter. **(A’’)** Complexity of IC (yellow) determined by the number of branches. (**B**) The schematic method used to slice IC into cross sections along its length, assisted by a Kimimaro skeleton, with a 3D reconstruction of IC **(B’)** and resulting 50-nm cross-sections (**B’’**). (Gray box indicates unusable sections). (**C**) Line plot depicting the major diameter (red) and normalized pixel intensity (blue) distributions along IC length. Gray zones indicate unusable sections shown in B’’ that were not considered for analysis. (**D**) Section from ssSEM volume (left) and 3D electron density reconstructions (right) showing the electron densities along an IC graphed in panel (C). (**E**) 3D reconstruction of **D** shown from two angles, with reconstructions of internal densities (midbody in cyan and microtubules in red), overlayed on a ssSEM slice containing the two GCs connected by IC.

To assess trends in the thickness (or diameter) of ICs, we sliced each IC at 50 nm intervals along their lengths to obtain a series of cross-sections (**Fig. 5B)**. To assist the determination of the axis for cross-sectioning, we skeletonized ICs using the Kimimaro algorithm (Silversmith et al., 2021) (**see supplementary methods**). We observed that cross-sections were largely circular, as evidenced by their elongation values **(see supplementary methods)**, ranging between 0.14 and 0.3 (average value 0.22). This circularity allowed us to compare cross-sectional diameter profiles of ICs, revealing a range of average diameters between 0.56 and 1.12 μm, in agreement with *in vitro* CB measurements (Mullins and McIntosh, 1982).

We then compared lengths and diameters of mitotic, migrating, and intermediate cell categories. IC lengths of mitotic cells ranged between 1.3 µm and 2.25 µm, compared to 1.85 µm and 2.6 µm in migrating cells **(Fig. 5A)**, indicating that connections can stretch longer in migrating cells than in mitotic ones. The length of connections between intermediate cells ranged from 1.6 µm and 3.25 µm.

Conversely, the connection diameter appeared larger in mitotic cells (from 0.85 to 1.082 µm) than in migrating cells (from 0.604 to 0.789 µm), with intermediate cells distributed mostly between mitotic and migrating cells (**Fig. 5A’**).

The morphological complexity of ICs was also analyzed. Unlike previously described membranous blebs adjacent to CBs (Sherman et al., 2016), several ICs had membranous protrusions (branches) **(Fig. S4B’, B”)** that shared their lumen with the shaft of the connection. Most ICs in mitotic cells had 8-10 branches, compared with 1 or 0 branches on the ICs of migrating GCs. Interestingly, the complexity of ICs on intermediate cells seems to be in between mitotic and migrating cells, with most having 2-5 branches (**Fig. 5A”**).

One of the most peculiar features observed within ICs were electron-dense regions, which we presumed to be microtubule bundles (Andrade et al., 2022; Zenker et al., 2017). We quantified the overall voxel intensity in each cross-section along ICs as a proxy for cytoskeletal filament density (**Fig. 5B-D, see supplementary methods)**. Profiles obtained from intensity-scaled electron micrographs showed that ICs contained dense regions that could correspond to MBs (**Fig. 5D**). Segmentation of darker voxels and 3D-navigation using the virtual reality software, DIVA (El Beheiry et al., 2020), illustrate these quantitative findings and provide a realistic representation of the cytoskeletal architecture within connections (**Fig. 5E)**.

We also identified several vesicular and membranous compartments within ICs. Segmentation of these compartments revealed volume and morphology heterogeneity, ranging from small spherical vesicles to large extended compartments (**Fig. S4A’, A”)**. The latter have not been found inside typical CBs. Overall, the cargos occupied a small fraction of IC volumes and were distributed throughout the bridge; however, the central dark intensities of ICs could also have obscured cargos in this area, meaning that the true cargo presence is greater than we could detect.

## Discussion

### Overview of the EM volume and segmentation

The complex properties of the adult mammalian brain arise from a finely orchestrated series of neurodevelopmental processes (Cajal, S.R., 1888). Abnormal laminar cytoarchitecture and cortical disorganization of neurons can lead to severe developmental disorders (Stoner et al., 2014).

In order to better understand the maturation process of dividing and migrating progenitor cells in the developing brain, we focused on cerebellar GCs located in the transient EGL for their ability to proliferate and migrate in a continuous, rolling fashion, and whose multiple stages of maturation into neurons can be studied using a 3D, ssSEM snapshot all at once (Stoodley, 2016; Wilson et al., 2019). Intriguingly, we found that both o- and iEGL contained GC pairs connected through ICs in the o- and iEGL.

Given that the oEGL at P7 harbors proliferating GCs, it is conceivable that some ICs correspond to CBs, however, the observation of GCs in prometaphase and telophase located in the iEGL suggests that GCs do not follow a deterministic maturation between EGL sub-layers, but rather, that they may be able to enter a mitotic state while in the iEGL. This discovery offers a new perspective to the study of the developing EGL and raises the question of whether dividing cells in different laminar layers have different precursors and/or whether they follow the same cell fate.

### IC-connected GCs exceed the expected number of GCs from prometaphase cells and show diverse morphology

Intriguingly, the total number of connected cells that the 11 cells in prometaphase could have generated (i.e. 22) did not match the number of IC-connected GCs (i.e. 60). If the 30 ICs were in fact CBs, an explanation could be that cytokinesis in GCs takes longer than mitosis. Another may be that at least some GCs deliberately retain their CBs in order to keep cells attached in a manner similar to IBs, stable CBs that form as a result of incomplete cytokinesis in spermatogonial cells (Braun et al., 1989; Mathieu et al., 2022). IBs could serve as a scaffold to transport key proteins to control cell polarization (Zenker et al., 2017), or synchronize future divisions (Espinosa and Luo, 2008). The observation of different cargoes, including membranous compartments and mitochondria, within ICs might support this hypothesis, as persistent IBs represent a mechanism of intercellular transport distinct from conventional vesicle trafficking in MBs of CBs.

Given the fixed-time-point constraint of our dataset, confirming the source of the unequal ratio of prometaphase and IC-connected cells is challenging. However, the fact that we observed neither cells interconnected as syncytia (i.e. more than 2 connected cells in a row) nor microtubule-bare ICs, both of which are features observed in IBs (Haglund et al., 2011; Mathieu et al., 2022), suggests that many of the ICs we observed differ from IBs previously described in gonad maturation (Haglund et al., 2011).

Our 3D-reconstructions revealed three specific features that differentiate ICs from classical CBs, each of which could give clues about the nature of ICs. First, IC-connected GCs are connected at locations other than their somas, may indicate a time-lag that gave CBs the opportunity to shift in position **(Fig. 3A)**. Second, ICs were wrapped by membranous, lamellipodia-like sheets that stemmed from both connected GCs **(Fig. 3B)**. Enveloping sheets could potentially play a role in protecting membrane remodeling and thereby in stabilizing the bridge. Alternatively, extended sheets could be in the process of engulfing the MB, which was proposed to be an intracellular signaling organelle that regulates cell proliferation (Peterman et al., 2019). Third, some IC-connected GCs in the iEGL have long (up to 30 μm) protrusions, suggesting that ICs (if they originate during cell division) may stay connected for a considerably long time after mitosis ends **(Fig. 3C, C’)**. GC protrusions have been considered to be precursors of parallel fibers, which immature GCs extend into the ML as they migrate down to the inner granule layer (Legué et al., 2015). This interpretation would suggest that our connected cells with protrusions are in the process of migrating out of the EGL.

### IC connected GCs-are in different stages: Mitotic, Intermediate, Migrating cells

The above observations indicate that at least some of these connected GCs exhibit characteristics unexpected for conventional mitotic cells. We grouped connected GCs into three categories, mitotic, intermediate, and migrating cells, using several cytoarchitectural features (**Fig. 4**). Based on this categorization, several inferences can be drawn about the behavior of each type of IC-connected GC.

All migrating IC-connected GCs have processes whose orientation follows that of parallel fibers, which indicate tangential migration, but some of them also show protrusions perpendicular to parallel fibers and their centrosome is located close to these processes. The centrosomes in the migrating cells are observed at different locations (in the soma or near the protrusion) which may suggest different stages of migration.

The connections between migrating cells could be remnants of mitosis or could be formed *de novo*. Dubois et al. (Dubois et al., 2021) described that the centrosome was implicated in Tunneling Nanotube (TNT) formation and that the Golgi complex was positioned in front of the connection, with the centrosome behind it. In addition, the centrosome has also been reported to be oriented towards the TNT. In migrating, IC-connected GCs, we observed this specific arrangement of the centrosome and the Golgi complex with respect to the connection for 7 of 12 GCs. In 4 of these, both the connected cells presented one centriole pointing towards the connection (**Fig. 4E**).

Intermediate GCs could then be considered as being chronologically located between mitosis and migration, following the hypothesis of a continuum of events represented by the presence of mitotic, intermediate and migrating cells. Such intermediate cells have been described as arising from asymmetric division and expressing markers of GC progenitors (MATH1), and early post-mitotic and tangentially migrating cells (DCX) (Yang et al., 2015). Furthermore, GCs in the developing chick cerebellum have been shown to grow and retract protrusions in between proliferative events, which could indicate that protrusion extension in GCs is not directly correlated with cell fate (Hanzel et al., 2019).

### Morphology of connections correlates with the category of connected GCs

The profiles of pixel intensity in the electron micrographs along connections provided an arbitrary proxy of the microtubule-like filaments running within ICs **(Fig. 5D)**. Such bundles of microtubules are well-known in cytokinetic bridges, and they form the midbody in the CB, along with several other proteins and vesicular cargoes. Interestingly we also observed various vesicular bodies forming cargo inside ICs **(Fig. S4A’, A”)**, which corresponds well with the vesicles shuttled by the microtubules within CBs for the completion of abscission in dividing cells (Skop et al., 2004). Our observation of a mitochondrion inside one of the ICs is perhaps indicative of their dissimilarity from conventional CBs.

Furthermore, the trends in length, diameter and complexity may indicate the changes in the morphology of these supposedly persistent CBs as the dividing cells progress towards migration **(Fig. 5)**. Indeed, we can imagine that it would be easier for two cells to migrate together, being connected by longer, thinner, and less complex structures, notably in a compact tissue environment.

### Known duration for protrusion growth largely exceeds the time required for mitosis

Finally, our IC connected GCs have long lamellipodial protrusions ranging up to 31.15 μm with an average length of 9.5 μm. The time lapse imaging performed by Hanzel et al. (Hanzel et al., 2019) in the developing chick cerebellum suggests that a protrusion of 29.14 μm would take ∼220 minutes to grow. At this speed, the longest protrusion of 31.15 μm in our connected GCs would require ∼235 minutes, while an average 9.5 μm long protrusion would take ∼72 minutes to grow. These durations are 3 to 9 times longer than the average time of mitosis, ∼25 minutes (Contestabile et al., 2009; Fujita, 1967), respectively. These results make us rethink our vision of individual cells undergoing well separated metabolic events.

## Conclusion/Speculations

While the limitations of our ssSEM-based approach do not allow us to explore the dynamics and biogenesis of cytoplasmic ICs between GCs, this technique provides an ultra-high-resolution view of the EGL of a developing mouse cerebellum. Detailed morphological characterization of these GCs reveals that their development in the EGL is more diverse and complex than conventional descriptions indicate. The wide range of features exhibited by ICs highlights the morphological diversity across connections and connected GCs. Our analysis indicates convincing signs of mitotic, migrating, and intermediate stages of connected cells, as evidenced by the presence and arrangement of organelles such as the Golgi complex, centrosomes, cilia, and cellular protrusions. ICs may be mitotic in origin, owing to their characteristics, such as microtubule bundles, presence of midbody-like densities, and dimensions similar to CBs. The hypothesis that ICs originate from mitosis suggests their persistence until the maturation of protrusions. Our rate estimations of protrusion growth suggest that GCs remain connected for several hours after mitosis. As such, irrespective of their origin, the observations of cells with long protrusions being connected may indicate that they are connected for purposes such as transfer of vesicles, mitochondria, and molecules between cells, as has been demonstrated in case of tunneling nanotubes (Gousset and Zurzolo, 2009; Rustom, 2004; Zurzolo, 2021). Recent results from Dubois and colleagues (Dubois et al., 2021) might also suggest that in some connected GCs, the centrosome close to the opening of the IC may be involved in their translocation through these connections or help in the formation of the de novo ICs. Overall, our results open several avenues of research to investigate the origin and function of ICs between developing GCs. Future research in living tissue may be required to diagnose the mechanisms underlying their formation and physiological function during cerebellar development, as well as assessing whether this phenomenon is widely present and relevant for CNS development.

## Acknowledgments

We thank members of the Lichtman lab (D.Berger, N. Dhanyasi, X. Lu, and X. Wang) for fruitful discussions; Jean Livet for critical discussions; all members of the Zurzolo lab for discussions and support, especially M. Rakotobe for critical discussions on cerebellum biology; W. Silversmith for early help during our initial skeletonizations; C. Godard for visualization of electron density using DIVA; data annotators: Ö. Özen, M, M. Trottmann, O. Mulhern, N. Borak, M. Bogan, A. Mancevski, R. McLellan, E. Sun, A. Atkin, S. O’Shea, N. Sullivan, A. Vaca, C. Vargas, D. Pacheco, A.P. Lastra, P. Chavan, C. Beltran, J. Mamani, A. Flores, A. Meruvia, F. Osalinas, T. Chrysostomou.

## Funding

Inception program (Investissement d’Avenir grant ANR-16-CONV-0005)

ANR-17-CONV-0005 Projet Q-life

Cancéropôle Ile-de-France 2018-1-PL BIO-03-IP-1

LEGS M. MICHEL

Equipe FRM

Third edition of the Big Brain Theory Call

Internal seed grant from IP to C. Zurzolo

U19 NS104653, U24 NS109102-01 and P50MH094271 Conte Center to J. W. Lichtman

C. Zurzolo was supported by the Radcliffe Institute for advanced study at Harvard (Fellowship program 2018-2019).

DCC was supported by the PPU program at IP.

## Author contributions

Conceptualization: CZ, JL, DCC

Dataset: AW

Methodology: AW, DCC, HK

Training of annotators: DCC

Software: HK

Investigation: DCC, HK, OCM

Visualization: DCC, HK, ND

Funding acquisition: CZ, DCC

Project administration: CZ

Supervision: DCC, CZ

Writing – original draft: DCC, HK

Writing – review & editing: CZ, DCC, HK, OCM, ND, AW, JL.

## Data and materials availability

Data can be provided upon request.

