## Supplemental Information for "3D reconstruction of the cerebellar germinal layer reveals intercytoplasmic connections between developing granule cells"

### Dataset

The serial-sectioning Scanning Electron Microscopy (ssSEM) dataset used in this study was obtained from the medial part of the cerebellar vermis in lobule VIII of a CD1 wild-type unsexed P7 mouse pup, previously published (available online at https://bossdb.org/project/wilson2019). The aligned and stitched image volume comprising 2514, 30-nm-thick sections cut in the sagittal plane, was imaged at 4-nm/px resolution. The volume was $1.7 \times 10^6$ $\mu m^3$ and contained regions from the cerebellar cortex: external granule layer (EGL), molecular layer (ML), Purkinje cell layer, and inner granule layer (IGL).

### Identification of cellular features in the ssSEM volume

Cerebellar granule cells: Identification of granule cell progenitors (GCPs) in the EGL was performed manually based on GCP location, size, morphology and nuclei features: the location of GCPs was determined by defining the boundaries of the EGL (see Reconstruction of the EGL|ML boundary below). At their widest point, the somas of GCPs had diameters of ~7 µm and exhibited a round-shaped body. GCPs contained nuclei that occupied most of the cytoplasm. Mitotic cells: GCPs in prophase were primarily distinguished by their disassembled nuclear envelope. The disassembled nuclear envelope was observed in the ssSEM volume as a fragmented membrane in the cell's cytoplasm. Cells in prophase also exhibited a spherical shape and lacked protrusions. GCPs in telophase were identified based on a wide, cleavage furrow that bridged the cytoplasm of two spherical-shaped cells; the nuclear envelope of both cells was already assembled. Intercellular connections (ICs) were identified based on a cylindrical tube shape connecting the cytoplasm of two cells; the presence of an electron-dense region resembling a midbody (MB), and microtubules emanating from the potential MB-ring into both cells. GCPs connected by ICs had fully formed nuclei. Tentative MBs were identified by a cluster of electron dense voxels inside the IC and predominantly between microtubule filaments on each side. Microtubule filaments were identified based on their electron density, straight and fine, (~25nm)-thickness. Cargoes within ICs were identified by membrane enclosure. Spherical cargoes included vesicles; non-spherical cargoes were labeled as membranous compartments. Mitochondria were distinguished by their shape and cristae. Protrusions extending out from the IC were classified as IC branches. Classification of branches was performed manually based on their 3D shape.

### Checks on dataset usability

Initial identification of cells in the EGL was performed manually via VAST (Volume Annotation and Segmentation Tool) (https://lichtman.rc.fas.harvard.edu/vast/) by annotating the nucleus of every cell, (or cell soma for cells with disassembled nuclei). Slight shifts in x & y during image acquisition of the dataset prevented exhaustive use of cells located at the top of the EGL, including all cells bordering the pia layer and cells on both side edges. To accurately estimate the number of cells in the volume, we employed the following rule: (1) classify cells as either (a) containing a fully visible nucleus in the volume (ITV), (b) partially visible out of the volume (OOV); (2) all cells ITV within a distance range of 7.5 µm from the center of the cell (the average thickness of a GCP), were flagged as likely-OOV. Of 2042 this method yielded an estimate of 932 GCPs.

### Tracing and rendering of ssSEM data

Regions of interest were manually traced with VAST. For rendering of all the traced images, 3D surface meshes of labeled objects were generated from VAST using VastTools (written in MATLAB) and imported into 3D Studio Max (Autodesk Inc.) to generate 3D renderings of all the traced objects.

### Cell level distribution analysis

The positions of mitotic and IC-connected GCPs along the radial axis in the EGL were assigned manually by counting GCPs, starting from GCPs bordering the ML and ending at GCPs nearing the pia. GCPs in levels 1-5 (1 being the layer closest to the ML) were classified as iEGL cells and GCPs in levels 6-10 (10 being the layer closest to pia) were classified as oEGL cells.

### CellWalker

The software CellWalker is scripted in Python (tested on version 3.x) and is available freely under the GNU General Public License (version 3.0) (https://github.com/utraf-pasteur-institute/CellWalker). Dependencies are described in the user manual (https://github.com/utraf-pasteur-institute/CellWalker). CellWalker supports image sequences of segmented microscopy images in PNG format. CellWalker exports various types of outputs depending upon the analysis performed, that include skeleton swc files, csv files for results of characterization and Wavefront

OBJ files for 3D visualization. These OBJ files can be directly imported in 3D rendering software Blender.

### Blender-python script for cross-sectioning the objects

This script is provided along with the CellWalker software. The usage is described in the manual (https://github.com/utraf-pasteur-institute/CellWalker/src/blender_python_scripts).

### Calculation of elongation of cross-sections

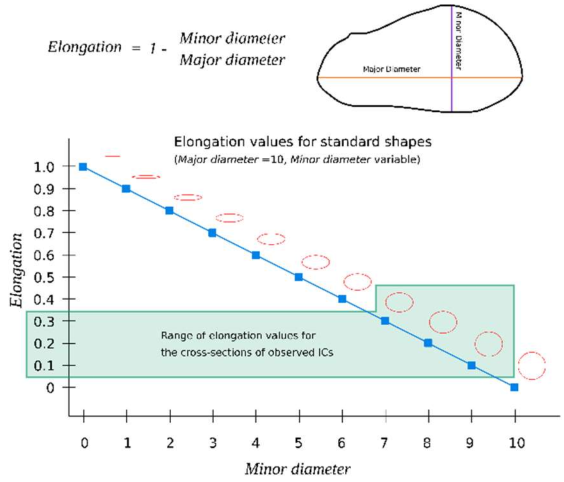


Elongation value was calculated for each cross-section along the connection to confirm the circularity of the cross-section.

Elongation of a 2D shape can be calculated as follows,

Elongation = 1 - (Minor diameter/(Major diameter)

The elongation range between 0 and 1, with values close to zero indicating circular shapes (See adjacent figure). The ICs in our data showed elongation values ranging between 0.14 and 0.3 (average value 0.22) indicating that the ICs were fairly circular in cross-section. Therefore we chose Major diameter as the diameter of the ICs.

### Cross-sectioning of IC along length

The ICs were cross-sectioned along their lengths using in the Blender software which works on the OBJ meshes exported from VAST or CellWalker. The cross-sectioning was assisted by a skeleton built using Kimimaro algorithm. The Kimimaro skeleton was built for an IC by loading its segmentation (exported from VAST) into CellWalker. CellWalker's skeletonization module helps applying the Kimimaro algorithm on selected segments in the loaded segmented image stack. The skeleton was visualized inside the CellWalker to manually identify the nodes on the skeleton that would form an axis approximately along the length of the connection. CellWalker's ability to export skeleton as OBJ file allowed us to load it into Blender and confirm that the selected nodes indeed formed an axis along the length of the IC. This axis (also called 'centerline') was also exported by CellWalker as an OBJ file. Next, the OBJ file of the IC (from VAST) and OBJ file for the centerline (from CellWalker) were loaded in Blender. A blender python script (provided in the Github repository above) was written to slice the mesh of the IC along the centerline at each point on the centerline (50 nm spacing decided by CellWalker). The blender python script also calculates the properties of the slices (or cross-sections) including major axis, minor axis and cross-section area (More features can be found in the script).

### Scaling and normalization of voxel intensities

All intensity values of the voxels were inverted, so that higher values correspond to darker regions in the ssSEM images. This design choice was made to facilitate the interpretation of our measurements of cytoskeletal density, since in this case the elements of interest are electron-dense.

In order to allow for comparison of cytoskeletal element density at bridges located in different regions of the ssSEM image volume, where pixel intensities may vary, we then min-max scaled intensities as follows.

$$I_{(i,scaled)} = \frac{I_i - I_{min}}{I_{max} - I_{min}}$$

where 'I' stands for intensity.

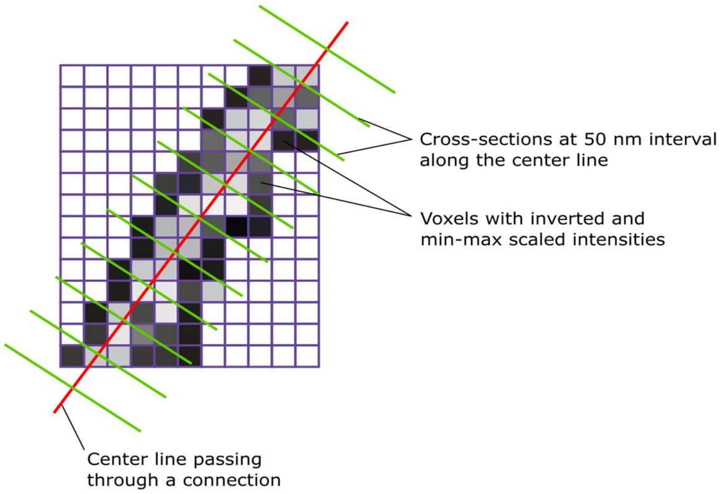


The sum of voxel intensities in a given slice was then obtained as a weighted average of the whole slice (see adjoining figure).

A voxel intensity profile along an IC is generated by generating a centerline (descript in previous section) along the connection and making cross-sections at 50 nm interval along the center line. The inverse voxel intensities are used to calculate a weighted average for each cross-section to generate the intensity profile along the connection. The profiles of this electron-density along all our ICs showed a peak within the IC corresponding to the region where darker microtubule-like bundles overlap (example shown in **Fig. 5C,D**). This peak also corresponds with a slightly increased diameter along the IC length (See **Fig. 5C**).

### Reconstruction of the EGL|ML boundary

The EGL|ML boundary is necessary as a reference for calculation of orientation of the connected cells. Conventionally and intuitively, the pia membrane is utilized for this purpose. However, in our ssSEM data, the pia is incomplete. The segmentation of the available traces of the pia indicated that it would be insufficient for calculating orientations of connected cells in the full volume. Therefore, we made use of the EGL|ML boundary as a proxy for the pia.

Manual identification of the ML through the entire data volume is an extremely time-consuming and intensive process. To make the process faster and reproducible, we designed a computational pipeline using deep learning classification followed by 3D point-cloud processing.

In short, the full pipeline can be described as follows. A Convolutional Neural Network (CNN) classifier was trained to detect cytological features to differentiate cells in the EGL from parallel fibers (PFs) in the ML and was then applied to uniformly sampled image tiles (300 x 300 pixel2) from the entire ssSEM volume **(Fig. S1)**. The training dataset consisted of randomly chosen and manually labelled EGL and ML tiles. In the inference phase the CNN was applied to uniformly sampled tiles from the full ssSEM volume. We used the coordinates of image tiles that were identified by the CNN as ML to create a point cloud and applied a ball-pivoting algorithm to construct a mesh demarcating the EGL|ML boundary. We then applied a cubic spline interpolation to remesh this course-grained surface and obtained a dense 3D surface (shown in light yellow in **Fig. 2A**).

The detailed procedure is as follows.

**Training data**

Four hundred tiles of size 300 x 300 pixel2 were randomly chosen from the ssSEM volume representing both EGL (200 tiles) and ML (200 tiles) classes. These tiles were subjected to augmentation in order to generate a larger training dataset as follows.

**Train/test split**

Each selected region was used to generate several augmented tiles. We split training images into train and test datasets prior to augmentation to ensure that augmented images from the same tile do not end up in both the train and test datasets. We split the 200 tiles of each EGL and ML class as 160 tiles in the train dataset and 40 tiles in the test dataset, i.e. 80:20 proportion.

**Data augmentation at source**

```
For each class (EGL and ML),
{
        For each tile
        {
                Apply shift in X and Y (Shift: 300 px at mip0 = 300/8 pixels at mip3)
                Original tile + 0 px shift
                Original tile + 300 px shift in X
                Original tile - 300 px shift in X
                Original tile + 300 px shift in Y
                Original tile - 300 px shift in Y
```

```
                    Resultant number of tiles = 5

                    For each shifted tile
                    {
                              Apply rotations: 0, 30, 45, 60, 90 degrees

                              Resultant number of tiles = 5
                    }
          }
}
```

Total number of augmented tiles:
For EGL class: 200 x 5 x 5 = 5000 (Train: 4000, Test: 1000)
For ML class: 200 x 5 x 5 = 5000 (Train: 4000, Test: 1000)

**Manual filtering**

These 5000 tiles belonging to each class are manually filtered to remove blank and partially out-of-volume tiles as well as ML tiles that contain portions of granule cells.

For EGL class after filtering: 4875 tiles (Train: 3900, Test: 975)

For ML class after filtering: 3400 tiles (Train: 2800, Test: 600)

The total number of manually filtered tiles is thus 8275 with 6700 (81%) tiles in the train dataset and 1575 (19%) tiles in the test dataset.

**Training/validation split**

Train dataset was split with a 70:30 proportion to obtain training and validation images before feeding to the CNN classifier.

**Stochastic augmentation (applied during the training process)**

Left-right flipping (mirror operation) with 0.5 probability

Random brightness (max_delta = 0.4)

Random contrast (lower limit 0.2, upper limit 0.6)

### CNN architecture

The CNN architecture was designed as illustrated in **Figure S1**. Our CNN has three convolutional layers with ReLU activation (kernel size 3x3) of sizes 32, 32 and 64 respectively, with a 2x2 max-pooling layer followed by each. The 'same' padding was used in all convolutional layers. The output of the convolutional block is flattened and a 50% dropout is applied as a measure against overfitting. After the 50% dropout layer, two dense layers with ReLU activation are inserted with 512 and 128 nodes respectively. Each dense layer is also followed by a batch normalization layer. In the end, a dense layer with a single node with a sigmoid activation is inserted to obtain the predicted value (0: EGL, 1: ML). We used an ADAM optimizer with a learning rate of 0.0001. The loss metric was binary cross-entropy.

The model with the smallest loss was chosen. Due to the clearly differentiable images of EGL and ML, the resulting model was extremely accurate, achieving F1-scores of 0.99897541 and 0.99833055 for EGL and ML respectively on the test data. The confusion matrix showed only 2 mis-classifications (out of 1575 total) where an ML image was classified as an EGL image **(Fig. S1)**.

### Inference (applying the classifier on the ssSEM volume to identify EGL and ML)

The tiles of size 300x300 pixels were sampled uniformly from the ssSEM volume with spacing of 100 px, 100 px in X & Y directions and 100 slices in the Z direction (at mip level 3), which correspond to approximately 3 um spacing. 15636 tiles were sampled such that they covered both EGL and ML. The trained CNN classifier was applied to these tiles and a cutoff of 0.9 was used to threshold the prediction score. This threshold ensured the correct identification of the ML tiles.

### Implementation of the CNN pipeline

The CNN pipeline was designed using Tensorflow-keras (version 2.4).

Training was performed on a Kaggle (https://www.kaggle.com/) kernel with Tesla P100 GPU. The training process was fast (9 s/epoch). Training batch size was set to 64 for optimal use of the GPU.

Inference was performed on a local computer (Dell Precision 3000 workstation with Intel Core i99900 processor and equipped with Nvidia Quadro RTX4000 GPU). Prediction on 15636 tiles sampled from the ssSEM took 6.8 seconds. The batch size was set to 64.

Our code is available as IPython notebooks on github (https://github.com/utraf-pasteur-institute/ssSEM-EGL-ML-analysis). Datasets can be made available upon request.

**Reconstruction of EGL|ML boundary surface using CNN predictions**

The center coordinates of the tiles identified as ML tiles by the CNN classifier were used to create a point-cloud using a python Open3D library. The normals were assigned to each point in the point-cloud such that they oriented towards the EGL. The ball-pivoting algorithm (as implemented in the Open3D library) was then applied to generate a mesh from the point-cloud. The assignment of normals facing in the direction of the EGL was necessary to ensure that the ball-pivoting algorithm identifies the required orientation in the point-cloud. The assignment of normals took care of the correct facing of the surface mesh such that it was created at the EGL|ML boundary.

This mesh was then exported as a .ply file (Stanford format) and loaded in the 3D rendering software Blender for visualization and manual fine-tuning. The fine-tuning was necessary because of the anomalies in the ssSEM data such as badly imaged regions (scratches, dark/light patches etc.), granule cells passing through the ML and also the nature of the EGL|ML boundary which is extremely uneven in certain regions. These anomalies led to the generation of holes in the EGL|ML boundary surface that needed manual attention. The fine-tuning step also involved deleting all other points in the point cloud that did not contribute to the EGL|ML boundary.

After closing the holes in the reconstructed EGL|ML boundary and removing unwanted points, we performed a remeshing operation using cubic-spline interpolation implemented in 3DSmax software. The resulting dense mesh (that contained approximately 54,000 points) representing the EGL|ML boundary was used for all further analysis.

The code is available as an IPython notebook on github (https://github.com/utraf-pasteur-institute/ssSEM-EGL-ML-analysis).

### Supplemental figures and legends

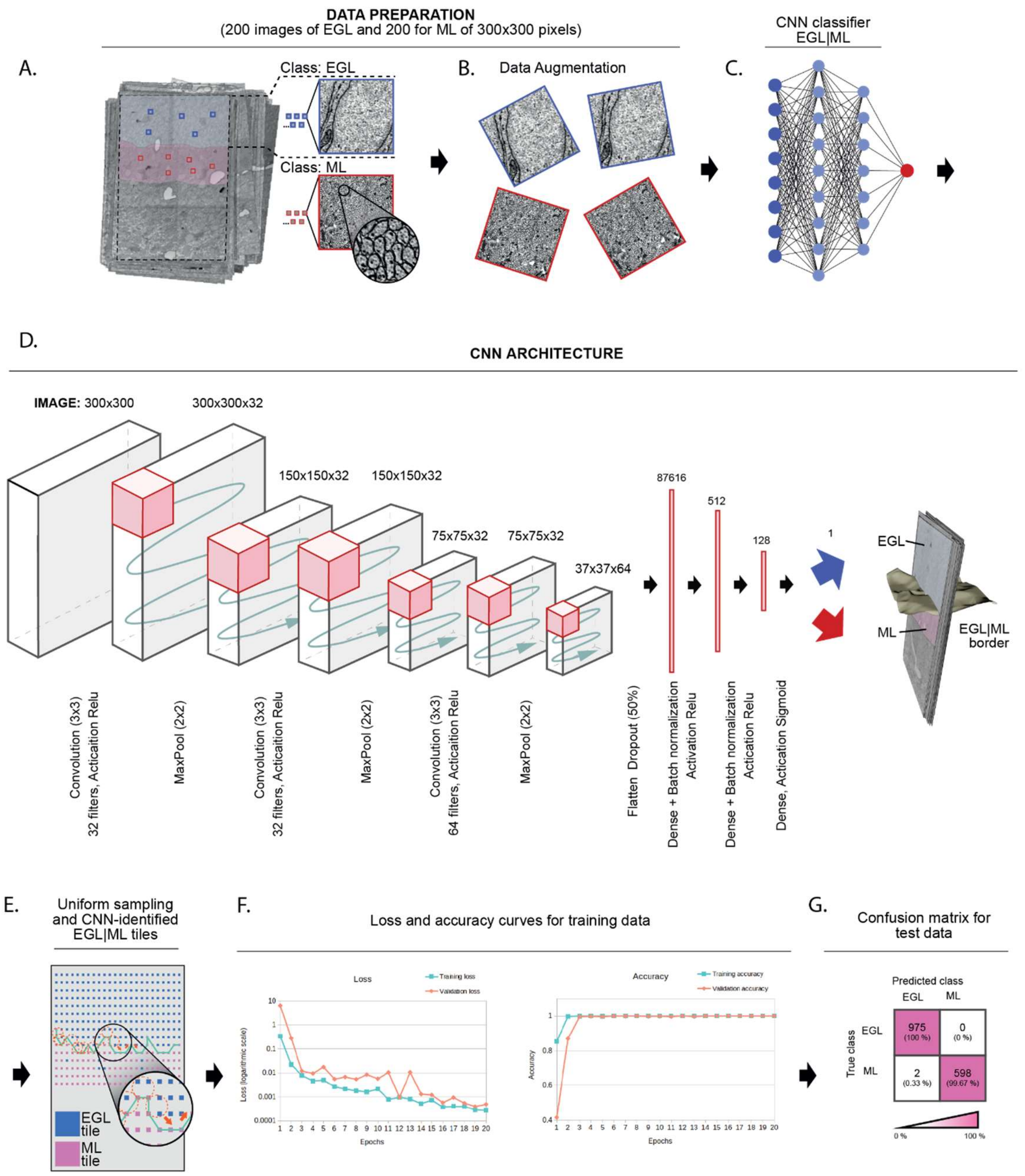


**Fig. S1. Overview of the training process of the convolutional neural network (CNN) designed to identify EGL|ML boundary (see supplementary methods for details)**

(**A**) Generation of training dataset: Manually chosen and labelled tiles for both EGL and ML classes from the ssSEM volume. (**B**) Data augmentation. (**C, D**) Architecture of the trained CNN classifier and the final result (**E**) Predicted labels (EGL/ML) for uniformly sampled tiles allowing the segregation of the two layers. (**F**) Training performance of the CNN- Loss and Accuracy curves saturating at close to zero and one respectively indicating satisfactory training. (**G**) Testing performance of the CNN- Confusion matrix showing near-perfect identification of EGL and ML classes in the test dataset.

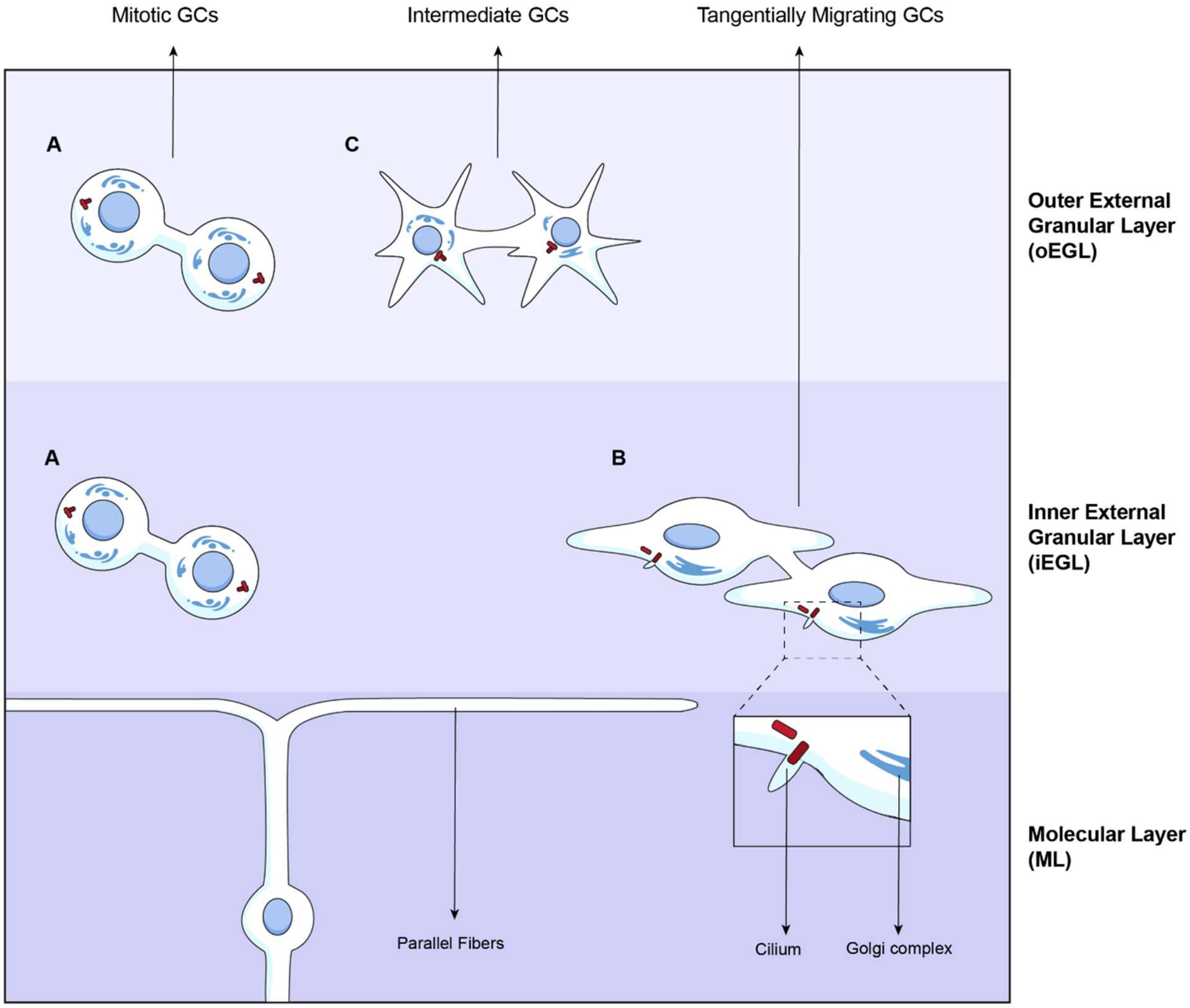


**Fig. S2. Categorization of GCs into three categories based on organelles and morphology of the cells, and positioning in the EGL of these cells shown in the coronal section.**

Connected GCs have been grouped in Mitotic, Intermediate and Migrating categories based on cell shape, number of Golgi complexes, presence of cilium, position of the centrosome in a cell and orientation of the protrusions of a cell when present (Categorization provided in **Fig. 4E**). **A)**

Connected mitotic GCs were mainly present in the oEGL, but some are also localized in the iEGL. They display more than two Golgi complexes dispersed in the cell, absence of a cilium, largely spheroid shape and centrosomes positioned at opposite poles of the cells. **B)** Connected tangentially migrating GCs are mainly localized in the iEGL. They harbor less than 2 Golgi complexes, have a cilium and show well-developed lamellipodia oriented parallel to the EGL|ML boundary and parallel fibers. **C)** Intermediate GCs were essentially observed in the oEGL. They share features of mitotic and migrating GCs. They show both filopodial and lamellipodial protrusions that are not specifically oriented in a particular direction, may have more than 2 Golgi complexes and a cilium, however the centrosomes are present in the soma indicating non-migratory behaviour.

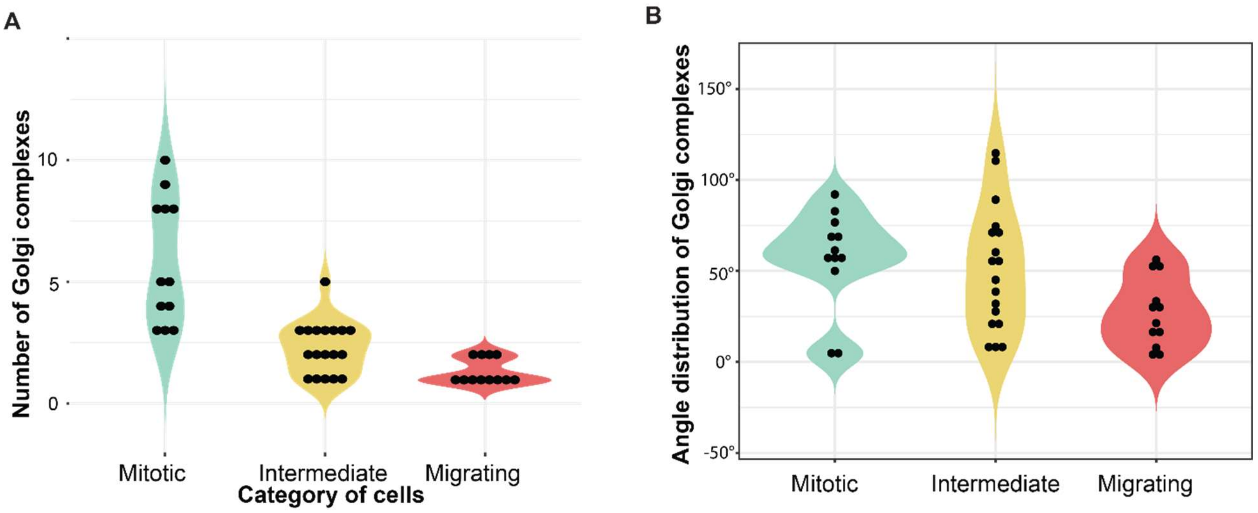


**Fig. S3. Morphometric characterization of IC-connected GCs among the different categories**

(**A**) Number of Golgi complexes observed in GCs of different categories (mitotic, intermediate, and migrating). In mitotic GCs, the Golgi complex is fragmented (3 to 10 individual Golgi complexes observed) compared to the Migrating GCs (1 to 2 individual Golgi complexes observed). (**B**) Angular distribution of Golgi complexes within GC. Angular distribution is calculated as the mean of the angles between the center of the nucleus of the GC and each voxel of the Golgi complexes in the cell. In mitotic GCs, Golgi complexes were homogeneously distributed in scattered clusters with most GCs displaying angular values above 50°, whereas most of the migrating GCs show angular values of Golgi complex distribution below 50° indicating well-formed Golgi complex.

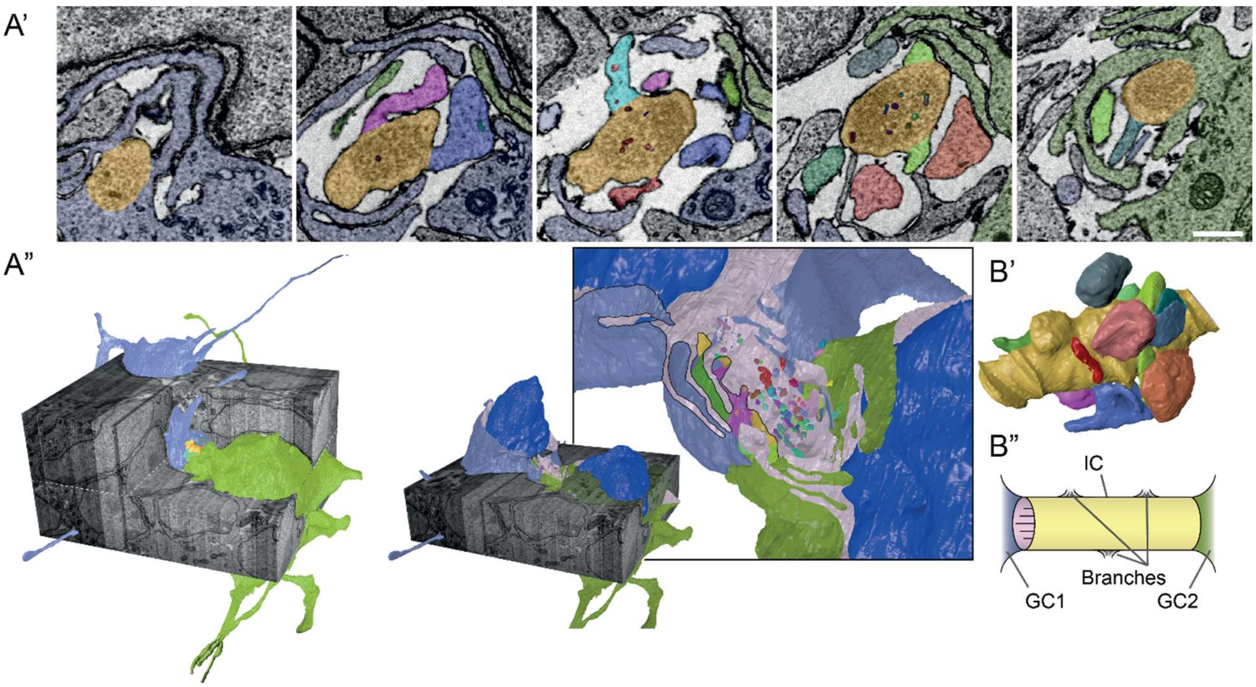


**Fig. S4.Subcellular elements associated with ICs**

(**A**) ssSEM micrographs (A') and 3D reconstructions (A'') showing GCs (blue and green) connected by IC (yellow) that contains numerous cargoes (vesicles, red arrow; long membranous compartments, cyan arrow, A''), and extending branch protrusions containing cargoes (magenta arrow, A'). (**B**) 3D reconstruction (B') and diagram (B'') of IC in A and its internal structure and contents, showing the branches that extend throughout.

**Movie S1.** 3D volume of a P7 mouse cerebellum imaged using ssSEM (provided as separate file)

**Movie S2.** GCs in EGL connected by ICs (provided as separate file)
